# Sex Differences in Cancer Functional Genomics: Gene Dependency and Drug Sensitivity

**DOI:** 10.1101/2025.02.05.636540

**Authors:** Nicole Zeltser, Chenghao Zhu, Jieun Oh, Constance H. Li, Paul C. Boutros

**Affiliations:** Department of Human Genetics, University of California, Los Angeles, CA, USA; Jonsson Comprehensive Cancer Center, University of California, Los Angeles, CA, USA; Department of Medical Biophysics, University of Toronto, Toronto, ON, Canada; Institute for Precision Health, University of California, Los Angeles, CA, USA; Broad Stem Cell Research Center, University of California, Los Angeles, CA, USA; Department of Urology, University of California, Los Angeles, CA, USA

## Abstract

Patient sex influences a wide range of cancer phenotypes, including prevalence, response to therapy and survival endpoints. Molecular sex differences across the central dogma have been identified that may drive these phenotypic differences. Despite a growing catalog of specific genomic, transcriptomic and proteomic sex differences in a range of cancer types, their functional consequences remain unclear. To assess how patient sex impacts cancer cell function, we evaluated 1,209 cell lines subjected to CRISPR knockout, RNAi knockdown or drug exposures. Despite limited statistical power, we identified pan- and per-cancer sex differences in gene essentiality in six sex-linked and fourteen autosomal genes, and in drug sensitivity for two compounds. These data expand our understanding of the propagation of sex-differing molecular features to functional outcomes, and their role in influencing patient phenotypes. They highlight the importance of considering sex-specific effects in mechanistic and functional studies.

## Introduction

Sex differences in cancer outcomes are well documented: incidence, mortality and response to treatment all differ between males and females^1–5^. These differences may in part be caused by differences in lifestyle, health-seeking behavior or other gendered characteristics. Other aspects of these differences appear to be a function of the differences in sex chromosomes and hormone levels (both during development and in adulthood). A broad range of correlative studies have evaluated molecular differences between male- and female-derived tumours, in hopes of explaining the differences in clinical outcomes. For example, tumours show sex differences in mutation patterns^6,7^, DNA methylation^8,9^, transcript abundance (Oh *et al*. in preparation), regulatory networks and protein abundance^7,8,10^ (Zhu *et al*. in preparation).

The wide range of molecular differences identified between tumours arising in men and those arising in women may in part explain the sex differences in clinical outcomes. However, the extent to which these tumour-intrinsic molecular differences have functional consequences remains underexplored. As a result, there remains a major gap between our detailed understanding of sex differences in clinical outcomes from both real-world evidence and trial analyses^1–5^, and between extensive descriptive characterization of somatic mutational sex differences in the tumour DNA.

Cancer cell lines provide a useful model system for large-scale functional genomic characterization, their molecular profiles being generally consistent with patient tumours^11^. The results of loss-of-function-based screens, implemented in pursuit of candidate drug targets, have high translational potential, and sex-specific effects may impact drug target discovery and validation efforts. Large-scale screening efforts have identified genomic, epigenomic and transcriptomic modulators of drug sensitivity and gene essentiality^11–15^, which in turn can be influenced by sex. While it is clear that sex chromosome dosage can influence pan-cancer gene essentiality^16^, it remains uncertain to what extent these observations are independent of the genetic technology used, whether they hold in individual cancer types, and whether they influence pharmacologic sensitivity.

To explore how sex differences in somatic mutations might relate to cancer cell function, we analyzed public Cancer Dependency Map (DepMap) project^17^ data from RNA interference (RNAi), CRISPR knockout and drug dependency screens, spanning 1,209 cancer cell lines^12,18–21^. Using a two-stage statistical approach^7,22,23^, we investigated sex differences in gene dependency and drug sensitivity on a pan-cancer and per-cancer scale, while adjusting for confounding variables. We identified 20 sex-linked and autosomal genes and two drugs with sex-differing sensitivity across a variety of cancers. These results encourage the consideration of the sex of tumour-derived cell line donors in any cell-line-based pharmacologic study and drug discovery effort.

## Results

### Sex differences in gene dependency and drug sensitivity

We evaluated gene dependency data for 19,720 unique autosomal and X-linked genes from CRISPR knock-out or RNAi knock-down screens and sensitivity to 4,739 unique drugs (PRISM1 and PRISM2 screens), derived from the Cancer Cell Line Encyclopedia (CCLE) DepMap database^17^ (**Supplementary Figure S1**). After excluding cell lines belonging to sex organ-specific cancer types and cancer types without at least one cell line of each sex, data were available for 1,209 cancer cell lines spanning 25 cancer types (**Supplementary Figure S2**). We used an established multi-stage statistical analysis approach^7,22,23^ (Oh *et al*. in preparation, Zhu *et al*. in preparation; see **Methods**). Univariable linear models with sex as a predictor were used to identify candidate sex-differing genes and drugs. Features that remained significant after multiple-testing correction were tested in multivariable models to adjust for cancer type-specific confounders. Separate analyses were performed on a pan-cancer and per-cancer level. For per-cancer analyses, after exclusions for a minimum sample size of three cell lines per sex category, 20 cancer types were included from the CRISPR dataset, 16 from RNAi, 14 from PRISM1 and 13 from PRISM2 (**Supplementary Figure S3**). LIHC (liver hepatocellular carcinoma), LUSC (lung squamous cell carcinoma), NKTL (T-cell lymphoma) and UVM (uveal melanoma), were only evaluated in pan-cancer analyses due to insufficient sample-size for per-cancer analyses. For multivariable analyses, after excluding cell lines without annotated covariates, three additional cancer types were excluded from per-cancer analyses in CRISPR and two from RNAi datasets. Coefficients and sample sizes from every combination of model (univariable or multivariable), feature and cancer type are reported in **Supplementary Tables 1-2**. Significant results from multivariable analysis are reported in **Supplementary Table 3**.

We first investigated sex-differing features on a pan-cancer scale, with sample sizes ranging from 299 (PRISM2, multivariable) to 584 (CRISPR, univariable) cell lines (**Supplementary Figures S2, S3**). After multivariable analysis, five pan-cancer sex-differing features were identified, all from the CRISPR dataset (**Figure 1A-B**). Dependency is measured on the CERES scale, where a lower score represents less cell proliferation and greater dependency on the knocked-out gene. All five genes showed a lower CERES gene effect score distribution in female cell lines compared to male cell lines (**Figure 1B**). All pan-cancer sex-differing genes were more essential to female cell lines (**Figure 1B**).

**Figure 1.**
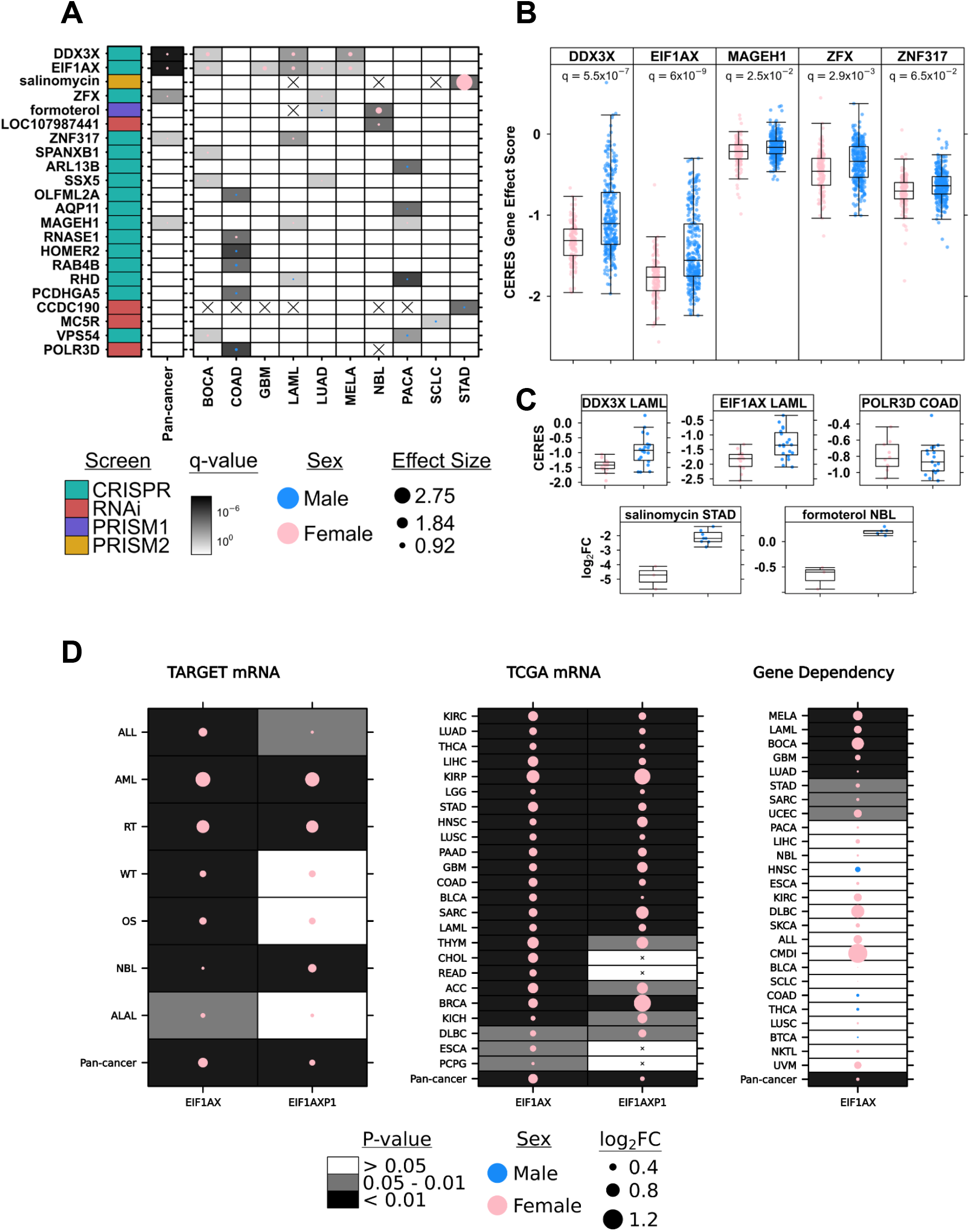
Sex difference in gene dependency, drug sensitivity and mRNA abundance. **A**. Autosomal & X-linked genes and drugs with significant sex differences in gene dependency or drug sensitivity on the pan-cancer level or in at least one cancer type based on multivariable linear regression (q-value < 0.1, MLR). Blank cells indicate cancer type and feature combinations with no significant difference by sex. Cells with crosses indicate cancer type and feature combinations that were excluded from analysis due to insufficient sample size. CRISPR = CRISPR knockout; RNAi = RNA interference knockdown; PRISM1 = primary drug screen; PRISM2 = secondary drug screen. **B**. CERES gene effect score from CRISPR knockout screening in the five genes from **A** with lowest q-value on the pan-cancer level. Lower score indicates greater sensitivity to gene knockout. **C**. Select cancer-specific effects from **A**. Gene sensitivity to CRISPR knockout is measured *via* CERES gene effect scores. Drug sensitivity is measured *via* log_2_ fold change of cell proliferation relative to negative controls. More negative log_2_ fold change indicates greater sensitivity to the screened drug. **D**. *EIF1AX* and its pseudogene, *EIF1AXP1*, have a higher mRNA abundance in females compared to males across all pediatric cancer types (TARGET, left) and adult cancer types (TCGA, middle) (p-value < 0.05, Mann-Whitney U test). *EIF1AX* also shows a consistent sex difference in gene dependency, favoring females, in most tested cancer types except HNSC, COAD, THCA and BTCA (right).

Next, we investigated functional genomic sex differences specific to individual cancer types. Per-cancer sample sizes ranged from our minimum threshold of three cell lines per sex (CRISPR analyses for THCA; thyroid adenocarcinoma) to a maximum of 85 (RNAi analysis of LUAD; lung adenocarcinoma). Sex-differing features (gene or drug) on a per-cancer level were found in ten cancer types, eight of which harbored more than one sex-differing feature, and five of which contained features from both directions of sex difference (**Figure 1A,C**). After multivariable analysis, significant sex differences on a per-cancer scale were identified in 22 features: 20 genes and two drugs. Of these, seven genes and one drug exhibited a significant sex difference in more than one individual cancer. All five genes identified in the pan-cancer analysis also differed in at least one individual cancer type (**Figures 1A-C, Supplementary Figures S4-9**). Among the 20 sex-differing genes detected in CRISPR knock-out or RNAi knock-down dependency datasets, none were independently replicated between the two perturbation technologies. Pan- and per-cancer p-value distributions indicate the potential presence of many additional associations at effect sizes below the detectable range of statistical power in this analysis (**Supplementary Figures S4-9**).

The X-linked gene *DDX3X*, homolog of Y-linked *DDX3Y*, has been implicated as a sex-differing tumor suppressor by higher loss of function mutation rates in male compared to female cancers^24^. In the CRISPR screen, *DDX3X* showed a greater dependency in female-derived cell lines, providing additional evidence of a sex-dependent functional effect in tumours. Sex differences were detected on a pan-cancer level as well as in BOCA (bone cancer), LAML (acute myeloid leukemia) and MELA (melanoma; q-value range [3.6 × 10^−3^,0.012]; effect size range [0.51,0.61]; **Figure 1C; Supplementary Table S3**). Another X-linked gene, *EIF1AX*, homolog of Y-linked *EIF1AY*, showed greater dependency in female cell lines on the pan-cancer level as well as in five specific cancer types: BOCA, GBM (glioblastoma), LAML, LUAD and MELA (q-value range [0.01,0.09]; effect size range [0.25, 0.64]; **Figure 1C; Supplementary Table S3**). Mutations and copy number alterations in *EIF1AX* have been shown to characterize uveal melanoma (UVM) subtypes, but there were not sufficient numbers of cell lines of both sexes to evaluate whether *EIFIAX* dependency differs by sex in UVM^25^. The gene *MAGEH1*, a member of the melanoma-associated antigen superfamily^26^, had marginally higher dependency in female-derived cell lines on the pan-cancer level (q-value = 0.025; effect size = 0.07). Two genes encoding zinc finger proteins, *ZFX* (X-linked) and *ZNF317* (autosomal), also had a higher female sex-bias on the pan-cancer level (q-value = 2.9×10^−3^ and 0.065; effect size = 0.13 and 0.09). Most genes with per-cancer sex differences in more than one individual cancer showed a bias in the female direction (female-derived tumours showing a greater dependency). A notable exception is *VPS54*, with opposite effects in BOCA and PACA (pancreatic cancer). COAD (colon adenocarcinoma) was the cancer with the greatest number of sex-differing genes, with all but one exhibiting greater dependency in male cell lines. All four X-linked genes identified with sex differences on a pan-cancer level have shown evidence of escape from X-inactivation^24^. Upon knockout, these genes may experience a greater relative reduction of expression in females than those normally expressed in only one X copy in males, leading to the observed impact on cell proliferation.

Two drugs, with no overlap between primary and secondary drug screens, were found to produce sensitivity that differed significantly by sex in cell lines from at least one cancer type. Drug sensitivity is reported as a log_2_ fold change (FC) in cell proliferation between treated cells and untreated controls. Salinomycin, a traditional anti-coccidial drug with anti-cancer activity in numerous cancer cell lines including gastric cancer ^27,28^, had higher sensitivity in female-derived STAD (stomach adenocarcinoma) cell lines at the highest screened concentration of 10 µM (q-value = 2.9×10^−4^; log_2_ FC = 2.75; **Figure 1A, C**). Another drug, formoterol, which has an anti-wasting effect in animal models with highly cachectic tumours^29^, had higher dependency in cell lines from male patients with LUAD (q-value = 0.047, log_2_ FC = −0.15) and female patients with NBL (neuroblastoma; q-value = 2.7×10^−4^; effect size = 0.98) at the only screened concentration of 2.5 µM (**Figure 1A, C**).

Overall, the influence of sex on functional outcomes in cancer cell lines is detected sporadically in our analysis, both on a pan-cancer and a cancer-specific level and in a technology-specific manner.

### Sex differences in EIF1AX and EIF1AXP1 replicates in mRNA abundance datasets

In an independent analysis of pediatric cancer transcriptomics in the TARGET database, the Chromosome 1 originating pseudogene *EIF1AXP1*, of X-linked *EIF1AX*, was found to have increased mRNA abundance in female pediatric patients with acute myeloid leukemia (abbreviated AML in TARGET and LAML by TCGA; Oh *et al*. in preparation). Given our similar finding of significant female bias in *EIF1AX* gene dependency in acute myeloid leukemia (LAML/AML) cell lines, we continued to investigate the pediatric and adult transcriptomic profiles of both genes across all available cancer types in the TARGET and TCGA databases. mRNA abundance for both genes differed significantly by sex in pediatric and adult tumours across the majority of tested cancer types (**Figure 1D**). Mann-Whitney U tests in TARGET and TCGA mRNA profiles showed a female bias in all tested cancer types for *EIF1AX* and 21 cancer types for *EIF1AXP1* (p-value < 0.05; **Figure 1D**). Mann-Whitney U tests in DepMap CRISPR knockout gene essentiality were also calculated for comparison (**Figure 1D)**. Cancer-specific sex differences in LAML/AML, GBM and LUAD replicated across DepMap gene dependency and TCGA mRNA datasets. LAML/AML was the only cancer type for which data was available in all three datasets, and sex differences were consistent across all three. Taken together, these results provide some evidence of a connection between transcriptomic profiles and functional genomic consequences in tumours.

## Discussion

Sex differences in cancer outcomes imply sex-differing functional consequences in tumour cell lines subjected to molecular perturbations. Our comprehensive analysis of the DepMap database identified a small number of such differences in gene essentiality and drug sensitivity screens. On a pan-cancer scale, we replicated prior findings of sex-chromosome dosage-dependent gene essentiality in the DepMap CRISPR knockout screen in *DDX3X, EIF1AX, ZFX and MAGEH1*^16^. Extending our analysis to additional screens and individual cancer types, we identified 10 cancers with sex-differing dependency on 16 additional genes and two drugs. These findings imply heterogeneous effects of patient sex on different tumour types, and that these effects persist in simple model systems. Sex-specific effects are likely also present in more complex models like organoids, an increasingly popular platform for high-throughput drug screening^30,31^. For effective generalization of results to both sexes, functional studies should include sufficient models to detect effects exclusive to each sex.

Gene essentiality quantifications from loss-of-function screens are contextualized by widespread mechanisms of genetic redundancy^32^. For example, the loss of a gene can be ameliorated by the expression of a functionally similar paralog with sufficient sequence divergence to avoid knockout. The paralogous regions of the X and Y chromosomes are excellent candidates for reciprocal buffering, and *DDX3X-DDX3Y, EIF1AX-EIF1AY* and *ZFX*-ZFY are experimentally validated in cell lines as functionally redundant paralog pairs^33^. In tumours, Y-paralog buffering can be undermined by the frequent, typically subclonal, loss of chromosome Y (LOY; 30% of all male TCGA tumours)^34^. LOY is estimated to occur in 30% of male cell lines in the DepMap database^16,33^, and LOY-mediated X-paralog essentiality in male cell lines has been demonstrated by others^16,33,34^. In our analysis, we expect the lack of Y-paralog redundancy in LOY male cell lines to have reduced differences in X-linked gene essentiality in comparisons of male-derived and female-derived cell lines. Despite this, we still detect significantly higher dependency in female cell lines on three X-linked genes with Y paralogs, suggesting these genes are robustly more essential in female-derived tumours. The essentiality of these and other genes differed by sex in some individual cancer types, but not others. Inter-cancer heterogeneity of LOY likely contributes to this discrepancy and reinforces the need for cancer-specific investigations.

Comparisons of functional studies are complicated by the differing mechanisms of screening technologies. For genes perturbed by both RNAi and CRISPR, significant differences did not replicate across screens, despite comparable sample sizes. This inconsistency may reflect technology-specific limitations: RNAi knockdown is more susceptible to off-target effects and inconsistent knockdown efficiency, while CRISPR knockout can be hampered by chromatin structure^35^. As CRISPR is generally more efficient, it is unsurprising that most gene dependency hits were detected *via* CRISPR screens. Nonetheless, the RNAi-specific findings highlight the value of incorporating multiple perturbation technologies in functional studies.

Mechanistic explanations for sex-differing functional outcomes are complicated by discordance of molecular findings across the central dogma and the complexities of sex chromosome genomics. Our autosomal gene findings have not been identified as sexually dimorphic in gene expression^7^(Oh *et al*. in preparation, Zhu *et al*. in preparation), consistent with the phenomenon of weak correlation between the genome & transcriptome and the proteome^36^. Molecular factors likely interact with a host of others to produce large sex differences in the complex phenotypes of cancer incidence and mortality. The majority of our X-linked gene findings also have not been recapitulated across the central dogma, due in part to the frequent exclusion of the sex chromosomes from molecular studies. Their unique structural features, including varying ploidy and repetitive or pseudo-autosomal regions, limit benchmarking and analysis efforts^37^ and create the potential for differential efficacy of specific molecular assays. As sequencing and bioinformatics technologies advance, more insights into the contribution of the sex chromosomes to cancer outcomes are anticipated.

Molecular sex differences in tumours are hypothesized to originate from a combination of genetic, endocrine and behavioral factors. These include the presence of differing sex chromosomes (X and Y), differing concentrations and secretion frequencies of gonadal and non-gonadal sex-linked hormones and differing rates of environmental exposure (*e*.*g*. smoking)^38–40^. The sex chromosomes differ conformationally between males and females, modified further by epigenomic phenomena such as incomplete X-inactivation, XX mosaicism and parental imprinting^38,39^. Conformational sex differences in autosomes have also been noted in mouse models^41,42^. Transcriptional sex differences begin at the earliest stages of embryonic development^38^. Several of these factors do not apply to immortalized cell lines, which are intrinsically limited model systems. In particular, cell lines lack interactions with cell-extrinsic factors known to mediate tumour biology, such as hormones^40^, cell-to-cell signaling^43,44^ and tissue architecture & cell morphology^45^. The nutrient composition of cell culture conditions does not perfectly replicate *in vivo* environments and variations in culture conditions exert selective pressures within cell line populations^46^. Cell-intrinsic factors in cell lines are also not completely representative of tumours. Prior work in DepMap cell lines has demonstrated substantial concordance between tumour and cell line molecular profiles^11,43^. However, cultured cancer cell populations are known to accumulate genomic variation, form subclone populations and exhibit gene expression patterns distinct from tumours^46,47^. Embryonic stem cell models exhibit an erosion of X-inactivation over time^48^. Historically, cell line to tumour concordance studies rarely report variation of concordance by epidemiologic characteristics of cell line donors. The stratification of cell-line-tumour concordance analyses by sex would provide additional context on the generalizability of our findings.

Another caveat to this study is the pervasive complexity of cancer, with myriad potential confounders to any analysis. To address this, we employed an established two-stage statistical approach, using univariable analyses to generate gene and drug candidates, followed by a multivariable model to control for confounders^7,22,23^. Unfortunately, cancer cell lines generally lack detailed clinico-epidemiologic annotation, making it difficult to adjust for features like race, smoking status and BMI. More consistent annotation of such features would expand opportunities for translational research.

Despite several limitations, our analysis sets a foundation for putative sex differences in tumour gene dependency and drug sensitivity. Given our findings of several sex-differing features, we suggest that sex should be considered in the interpretation and design of large pharmacologic and genetic screening studies.

## Acknowledgments

This work was supported by the NIH/NCI under awards P30CA016042, R01CA244729, U01CA214194 and U2CCA271894. NZ was supported by NHGRI Training Grant in Genomic Analysis and Interpretation T32HG002536 and by NCI training fellowship F31CA281168. CZ and JO were supported by the UCLA Jonsson Comprehensive Cancer Center fellowship program.

## Completing Interests

P.C.B. sits on the Scientific Advisory Boards of BioSymetrics Inc and Intersect Diagnostics Inc., and formerly sat on that of Sage Bionetworks.

## Author Contributions

**Initiated Study:** PCB, CZ, JO, NZ, CHL

**Data Analysis:** CZ, JO, NZ

**Supervised Research:** PCB, CZ, JO, NZ

**Wrote First Draft of Paper:** CZ, JO, NZ

**Approved Paper:** All authors

## Methods

### Gene dependency and drug sensitivity

#### Datasets

Four datasets from the DepMap Consortium were used to evaluate sex differences in cancer cell line gene dependencies: the CRISPR knockout genetic dependency screen, the RNA interference knockdown genetic dependency screen and the primary (PRISM1) and secondary (PRISM2) drug repurposing sensitivity screens^12,18–21^. PRISM screening is a high-throughput method for profiling the sensitivity of cancer cell lines to drugs, where the primary screen (PRISM1) identifies active compounds, and the secondary screen (PRISM2) validates their activity across a range of concentrations. Gene dependency and drug sensitivity matrices were downloaded from the DepMap data portal. CRISPR knockout screen data were downloaded from release 21Q1^18–20^, RNAi knockdown screen data from the DEMETER2 v6 data files^21^, primary drug screen data from PRISM release 19Q3^12^ and secondary screen data from PRISM release 19Q4^12^. The cancer type of each cell line was annotated by mapping the DepMap tumour designations for each cell line, consisting of the combination of primary disease and disease subtype, to the closest equivalent project codes as used by The Cancer Genome Atlas^49^ (**Supplementary Table S4**).

#### Cell line exclusion criteria

Cell line samples were excluded from all analyses if belonging to cell lines derived from sex organ-specific tumours (ovarian, uterine, cervical, prostate, testicular), belonging to cell lines of cancer types without at least one cell line of each sex or lacking an annotation of donor sex. Cell lines with missing information on the age of their donors were excluded from multivariable analyses. Cell lines belonging to cancer types which did not meet the minimum sample size threshold of three cell lines per sex category were excluded from per-cancer analyses.

#### Feature exclusion criteria

Genes on the Y chromosome in CRISPR and RNAi datasets were excluded. The PRISM2 secondary drug screen screens each unique drug at multiple concentrations. Each combination of drug and concentration was included and tested independently. DepMap data matrices report missing gene essentiality or drug sensitivity measurements for some combinations of cell lines and features (gene or drug). For each combination of feature and cancer type, if exclusion of cell lines with missing measurements resulted in fewer than three cell lines per sex category, the feature was omitted from analysis (coefficients reported as “NA” in **Supplementary Tables 2-3**).

#### Statistical Framework

Sex differences in gene dependency and drug sensitivity were evaluated using multiple stages of statistical testing. Each dataset was evaluated separately. First, for each feature (gene or drug), univariable linear models with sex as the predictor variable were applied at both the pan-cancer level and separately for each cancer type (per-cancer analyses). The False Discovery Rate (FDR) method was applied to correct for multiple testing separately on per-cancer and pan-cancer results. Next, the pan-cancer analysis in all features was repeated with multivariable linear regression using sex as the predictor and age and cancer type as covariates. Features with a q-value < 0.1 in pan-cancer univariable, pan-cancer multivariable or per-cancer univariable models were considered putatively associated with sex. For the final analysis step, only putative sex-differing features were included in per-cancer multivariable linear regressions, with sex as a predictor and age as a covariate. Per-cancer multivariable results were once more corrected for multiple testing using FDR. Features with a q-value < 0.1 in multivariable analyses were reported as having demonstrated a significant sex difference.

### EIF1AX and EIF1AXP1 transcriptomics comparison

The Cancer Genome Atlas (TCGA) mRNA TPM abundance matrices with 7,837 patients with 25 different cancer types were downloaded through the Genomic Data Commons (GDC) portal^50^ using TCGAbiolinks ^51–53^. The 25 analyzed cancer types were selected with the criteria of a minimum of 3 patients in either sex group. TPM values of EIF1AX and EIF1AXP1 were extracted from the abundance matrices and a Mann-Whitney U test was performed on each cancer type as well as between aggregated pan-cancer data. Therapeutically Applicable Research to Generate Effective Treatments (TARGET) initiative mRNA FPKM-UQ profiles were downloaded through the GDC data portal. Clinical data was available through the TARGET data matrix. All cancer types with available data were included in the analysis: acute lymphoblastic leukemia (ALL), acute myeloid leukemia (AML), neuroblastoma (NBL), osteosarcoma (OS), Wilms’ tumour (WT), rhabdoid tumour (RT) and acute leukemia of ambiguous lineage (ALAL). A Mann-Whitney U test was also applied to solely to *EIF1AX* gene dependency data, as *EIF1AXP1* was not measured in the DepMap dataset. All tests were two-sided and an alpha level of 0.05 was used to report statistical significance.

## Data availability

All DepMap data are available for download through the DepMap portal: https://depmap.org/portal. The results shown here are in part based upon data generated by the TCGA Research Network: https://www.cancer.gov/tcga. The pediatric results published here are in part based upon data generated by the Therapeutically Applicable Research to Generate Effective Treatments (TARGET) initiative, phs000218, managed by the NCI. The data used for this analysis are available at https://portal.gdc.cancer.gov/projects^50^, https://ocg.cancer.gov/programs/target/data-matrix and https://www.ncbi.nlm.nih.gov/projects/gap/cgi-bin/study.cgi?study_id=phs000218.v24.p8 (sub-studies phs000463, phs000464.v21.p8, phs000465.v21.p8, phs000466.v21.p8, phs000467.v21.p8, phs000468.v21.p8, phs000470.v21.p8, phs000471.v21.p8). Information about TARGET can be found at http://ocg.cancer.gov/programs/target.

## Code availability

All statistical analysis and data visualization were performed in the R statistical environment (v4.0.2). All visualization were performed using the BPG (v6.0.3)^54^. Code is available at https://github.com/uclahs-cds/project-CancerBiology-SexDifferencesv3PedProtFunc.

---

**Supplementary Figure S1. A**. Venn diagram of counts of unique autosomal & X-linked genes analyzed for sex differences in gene essentiality between CRISPR knockout and RNAi knockdown screens in the DepMap database. **B**. Venn diagram of counts of unique drugs analyzed for sex differences in drug sensitivity between PRISM1 primary screens (all drugs at the same concentration), and PRISM2 secondary screens (each drug was screened at eight concentrations, drugs screened multiple times are counted once in this figure).

**Supplementary Figure S2**. Summary of 1,209 cell lines in the DepMap database derived from cancer types included in pan-or per-cancer analyses of sex differences in gene essentiality and drug sensitivity. Cell lines belonging to sex-organ specific cancer types or cancer types without at least one cell line per sex category were excluded from all analyses and are not shown here. Central heatmap indicates included cell line availability by source screen dataset. Not all cell lines are screened in all datasets. Covariate bars (right) indicate cell line sex, cancer type, inclusion in pan-/per-cancer models and univariable/multivariable models. Top panel displays cell line totals by sex for each dataset.

**Supplementary Figure S3**. Counts of cell lines by cancer type included in per-cancer univariable (top) and per-cancer multivariable (bottom) analyses in each source screen dataset. Cancer types with fewer than three cell lines in each sex category are excluded.

**Supplementary Figure S4**. P-value distributions from univariable linear model analyses associating sex with CRISPR knockout induced gene dependency (top), RNA interference knockdown induced gene dependency (middle) and drug screen induced sensitivity (bottom) across pan-cancer analyses (left) and in each cancer individually (right). n = number of tests. Drugs screened at multiple concentrations were tested independently.

**Supplementary Figure S5**. P-value distributions from multivariable linear model analyses associating sex with CRISPR knockout induced gene dependency (top), RNA interference knockdown induced gene dependency (middle) and drug screen induced sensitivity (bottom). Pan-cancer analysis, adjusted for age and tumour type, performed on all genes and drugs passing inclusion criteria (left). Only genes and drugs with significant associations in univariable linear model analyses and pan-cancer multivariable linear model analysis were analyzed in each individual cancer, controlling for age (right). Features were excluded from per-cancer analyses if fewer than three cell lines per sex category were remaining after exclusion of missing covariate data. n = number of tests. Drugs screened at multiple concentrations were tested independently.

**Supplementary Figure S6**. P-value distributions from multivariable linear model analyses in each cancer type included in per-cancer analysis in the CRISPR knockout dataset. n = number of genes tested.

**Supplementary Figure S7**. P-value distributions from multivariable linear model analyses in each cancer type included in per-cancer analysis in the RNAi knockdown dataset. n = number of genes tested.

**Supplementary Figure S8**. P-value distributions from multivariable linear model analyses in each cancer type included in per-cancer analysis in the PRISM1 primary drug screen dataset. n = number of drugs tested.

**Supplementary Figure S9**. P-value distributions from multivariable linear model analyses in each cancer type included in per-cancer analysis in the PRISM2 secondary drug screen dataset. n = number of drugs tested. Drugs screened at multiple concentrations were tested independently.

**Supplementary Table 1**. Statistical coefficients of gene dependency and drug sensitivity using univariable analysis in cancer cell lines.

**Supplementary Table 2**. Statistical coefficients of gene dependency and drug sensitivity using multivariable linear regression in cancer cell lines.

**Supplementary Table 3:** Statistical coefficients of gene dependency and drug sensitivity using multivariable linear regression in cancer cell lines in features with a false discovery rate lower than 0.1.

**Supplementary Table 4**. Mapping of cancer types codes assigned to CCLE cancer descriptions.

