## Supplementary Figures S1-S9 for "Sex Differences in Cancer Functional Genomics: Gene Dependency and Drug Sensitivity"

#### Supplementary Figure S1

**A**

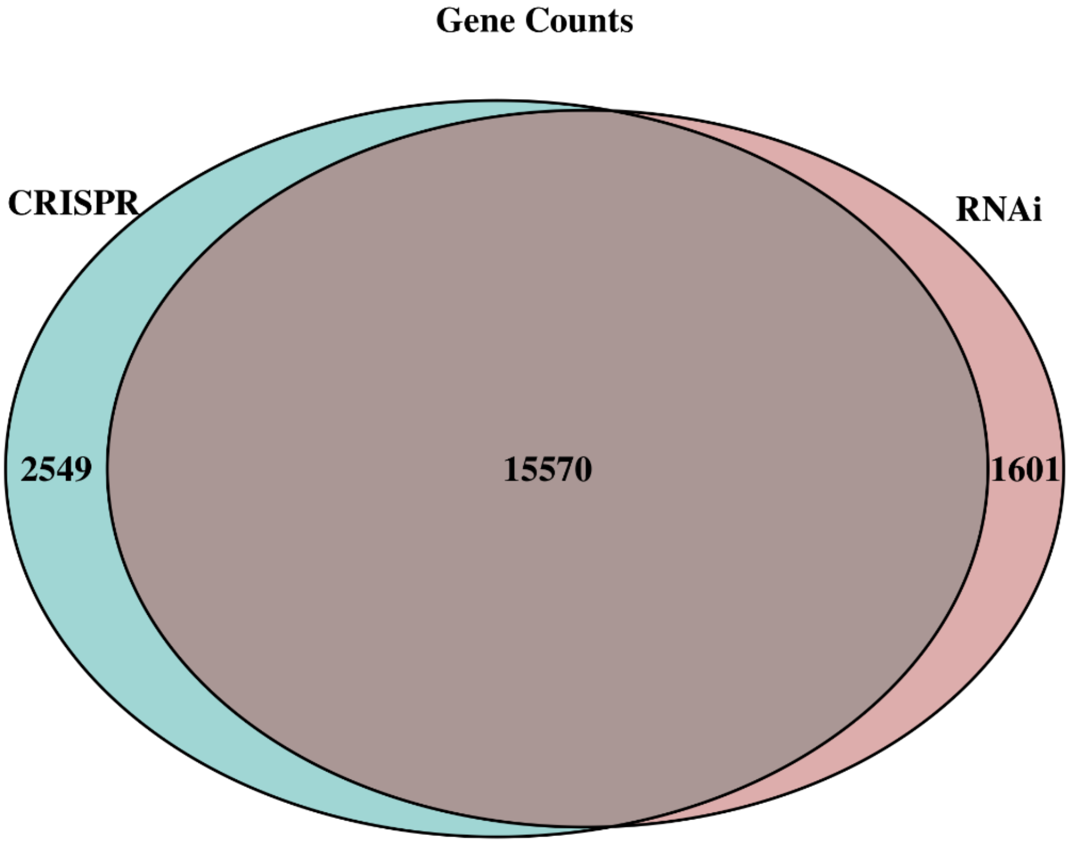


**B**

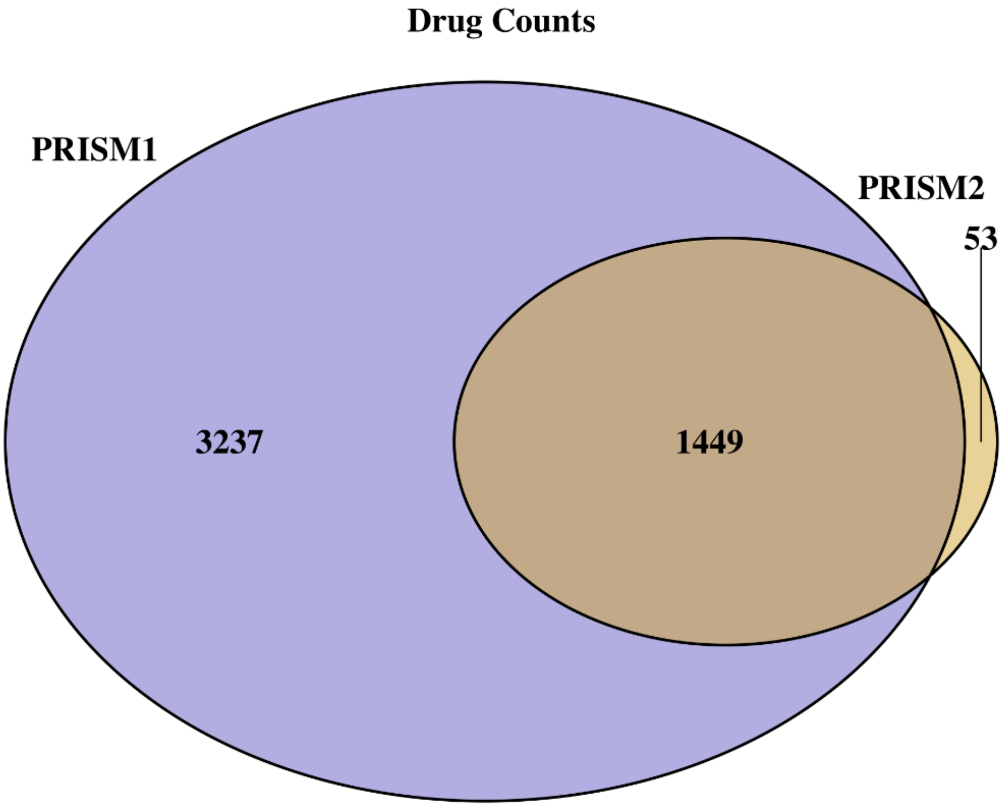

### Supplementary Figure S2

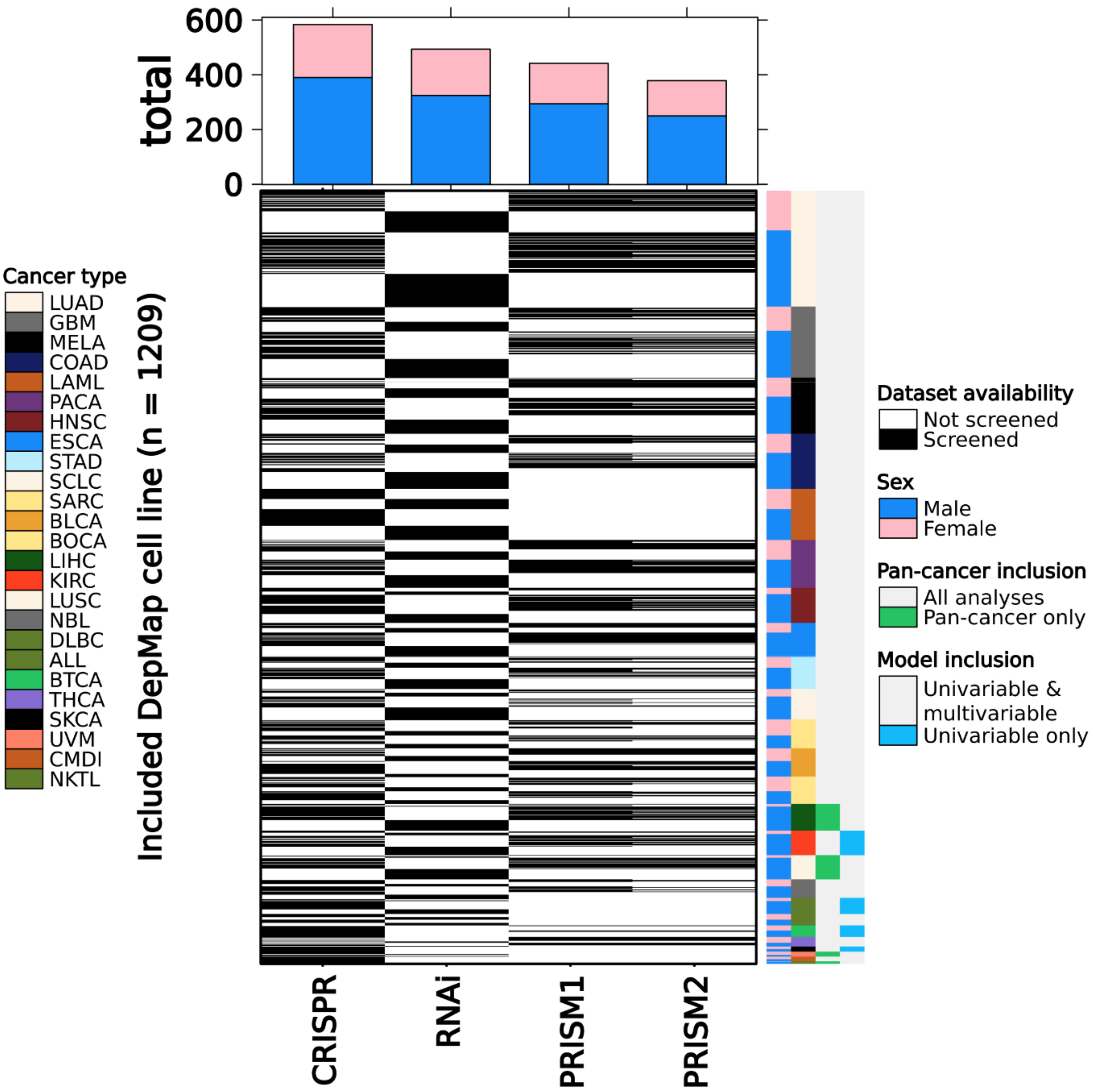

### Supplementary Figure S3

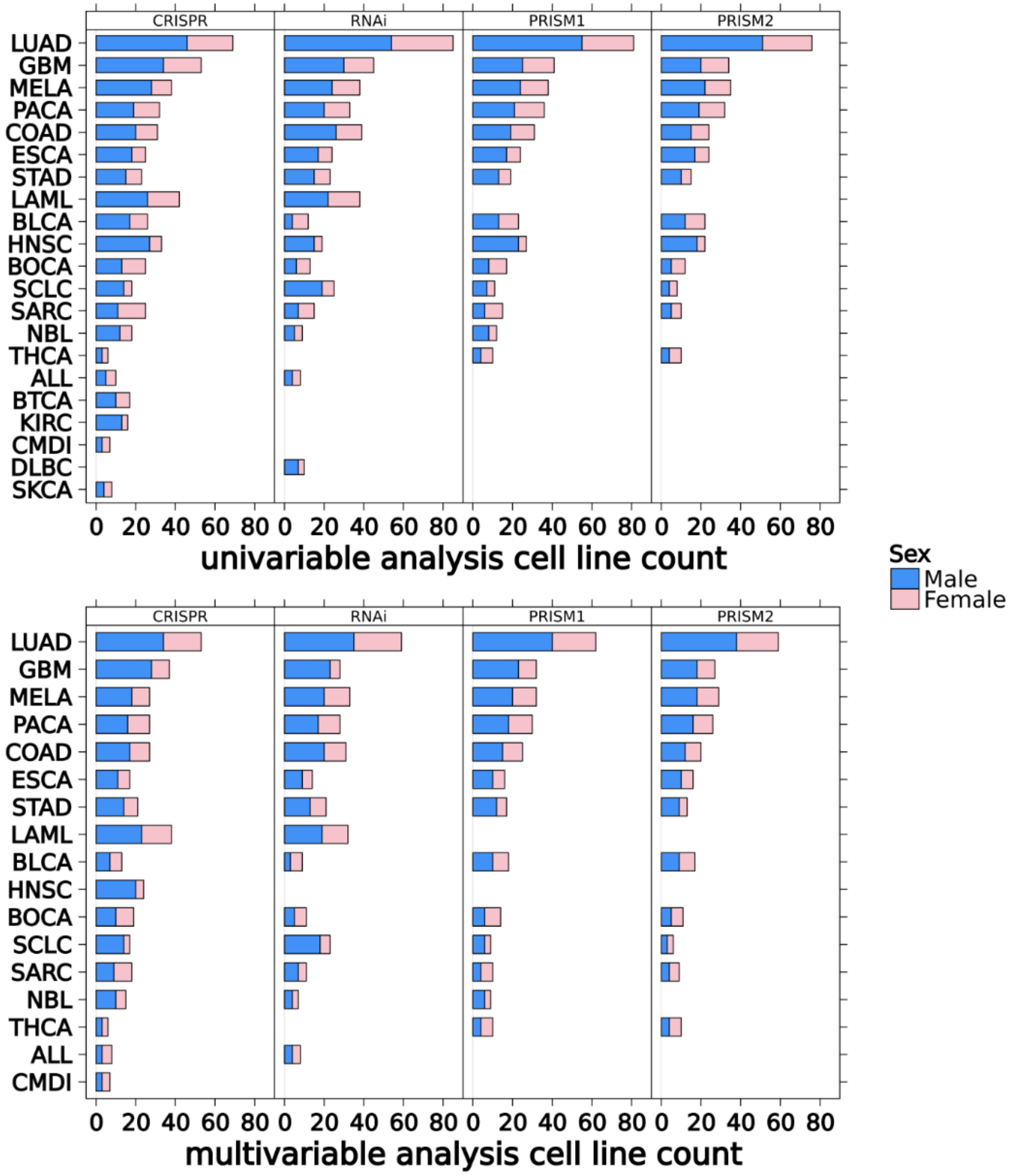

**Supplementary Figure S4**

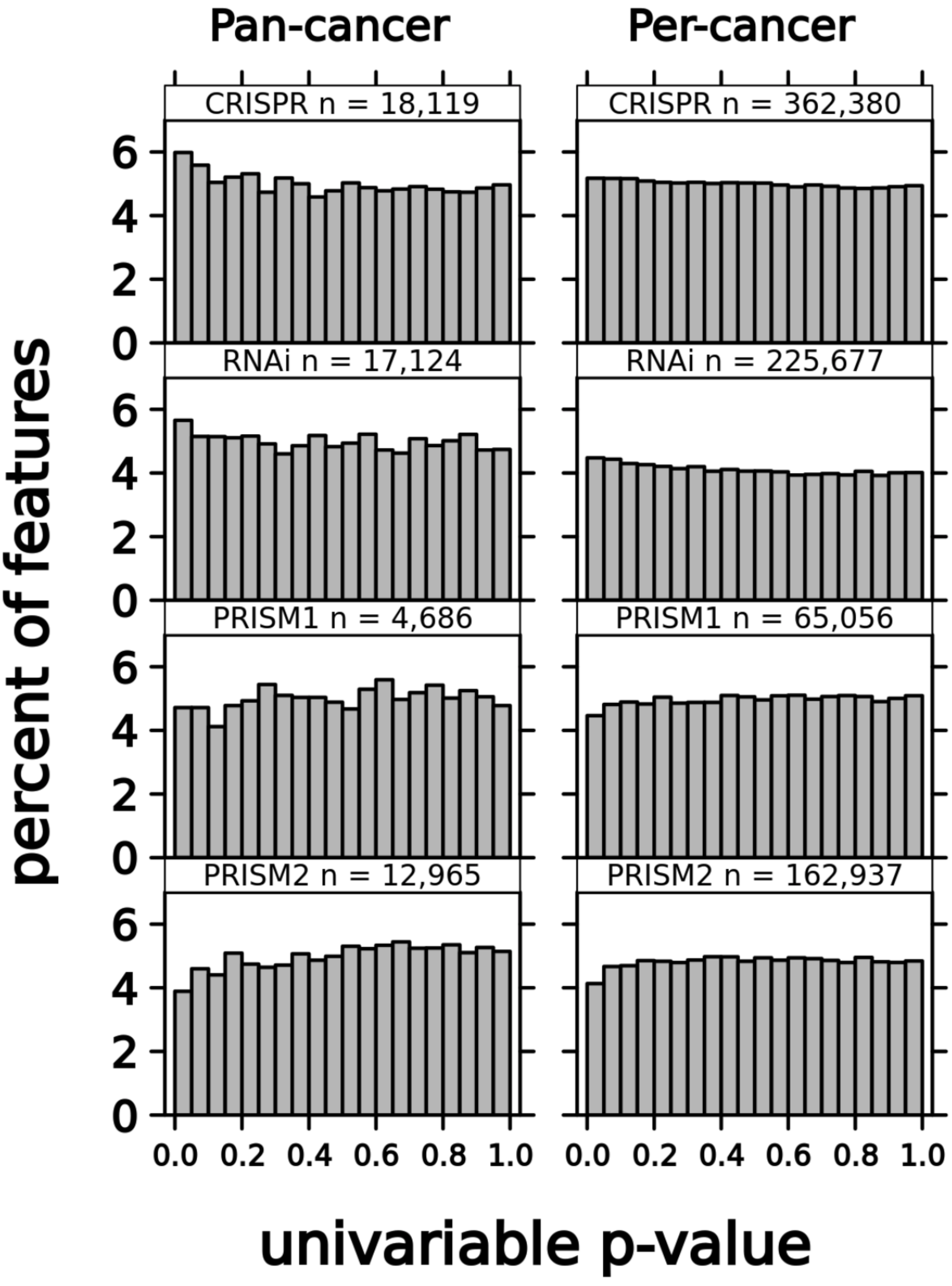

Supplementary Figure S5

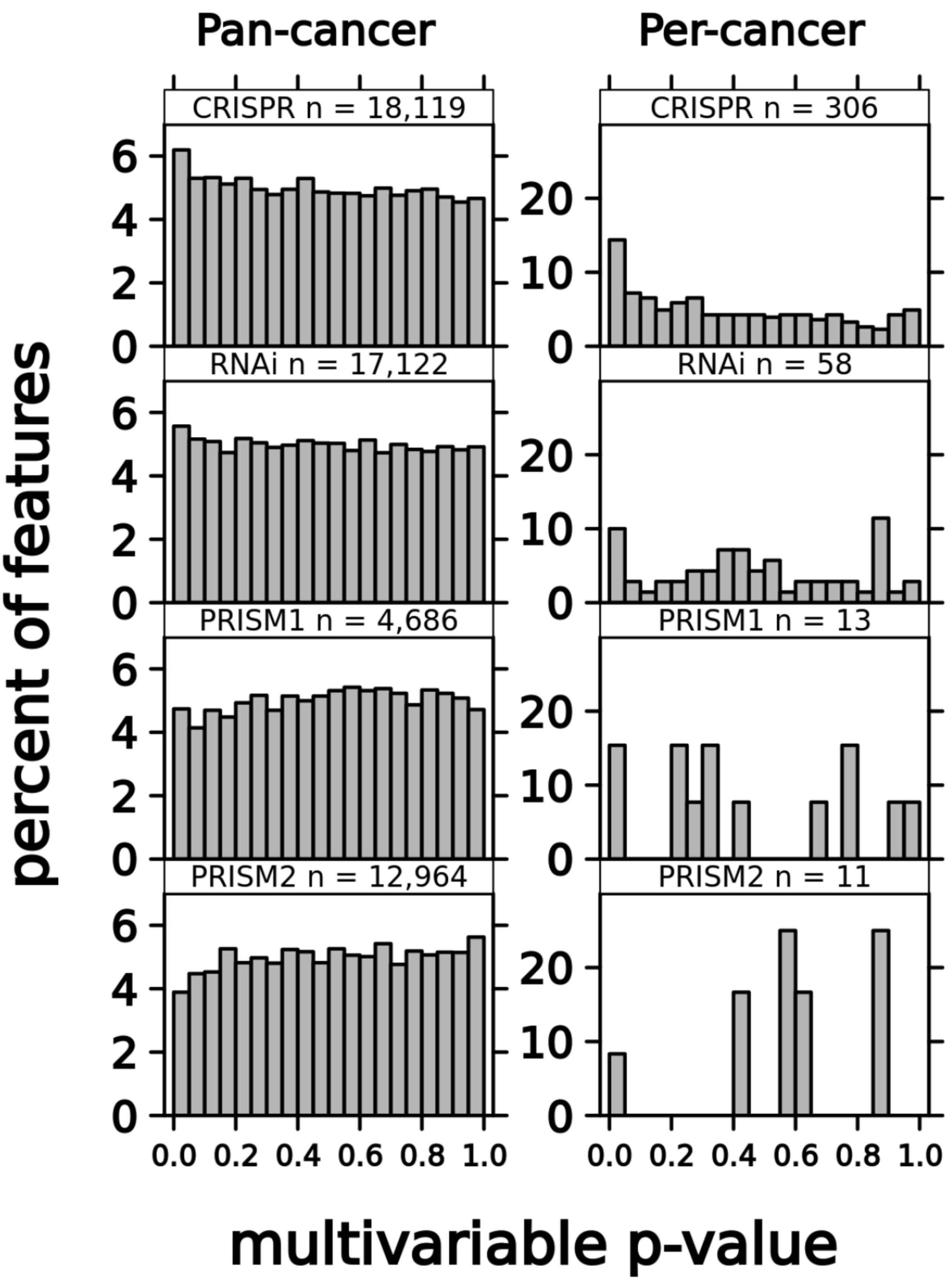

#### Supplementary Figure S6

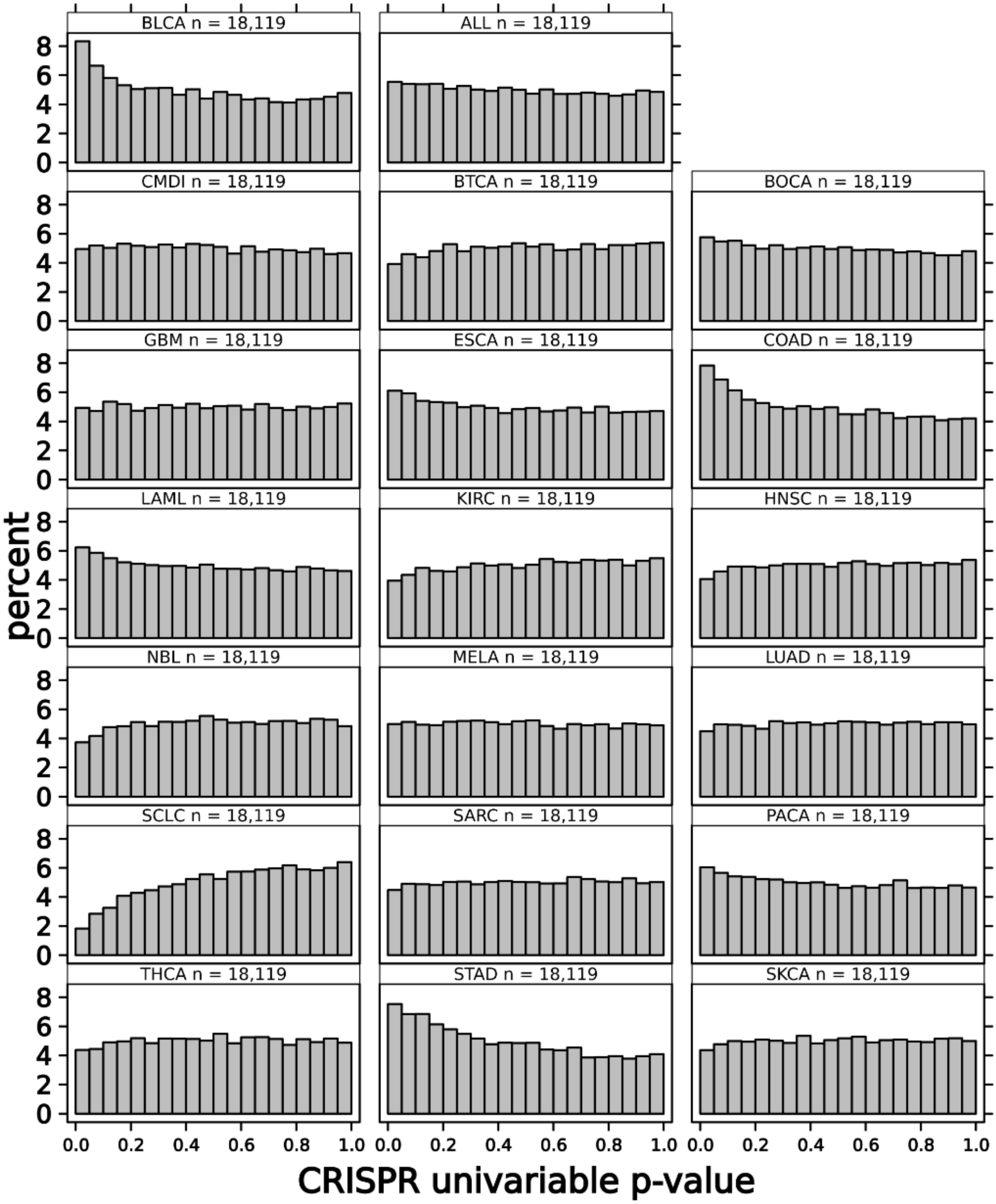

### Supplementary Figure S7

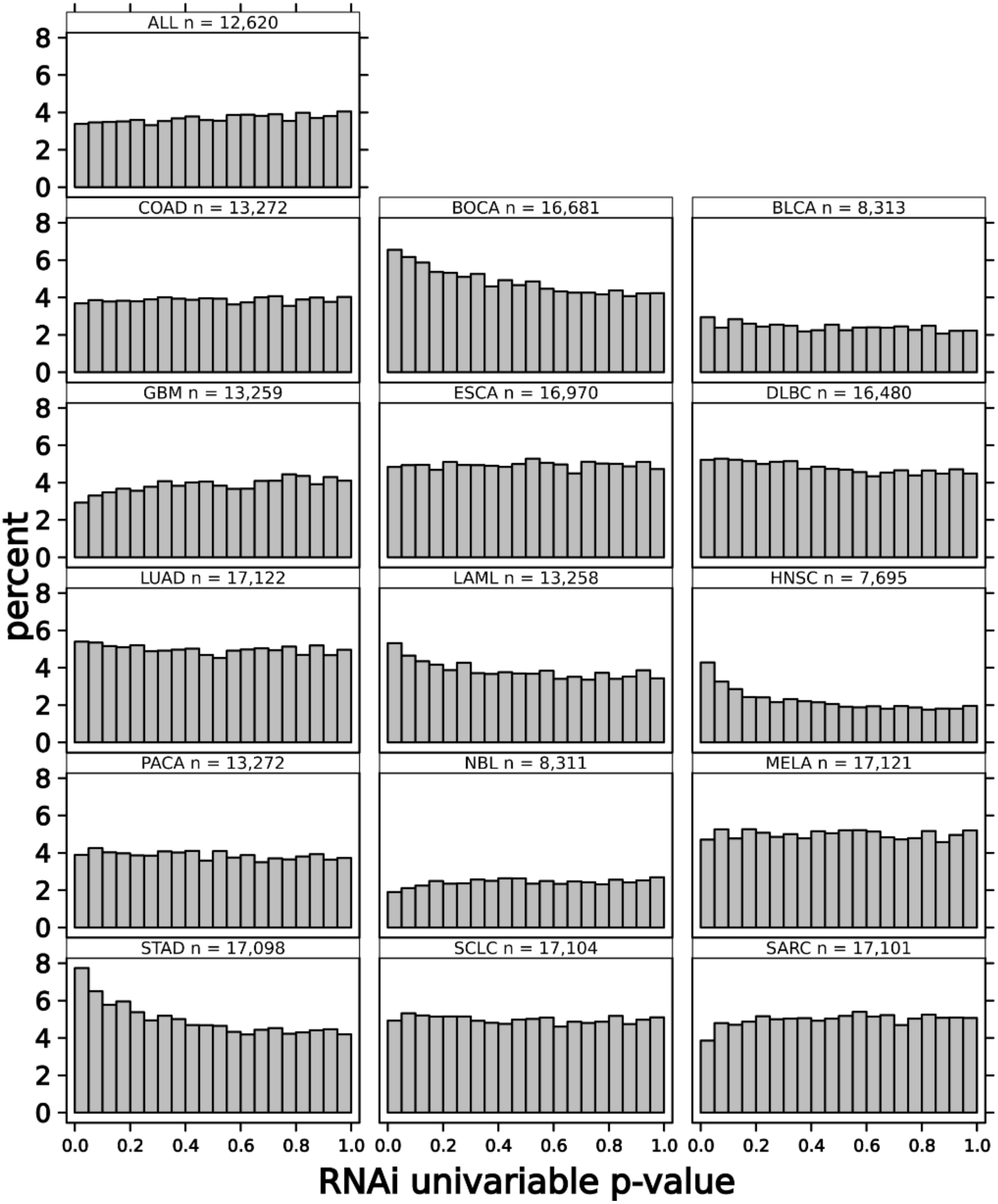

**Supplementary Figure S8**

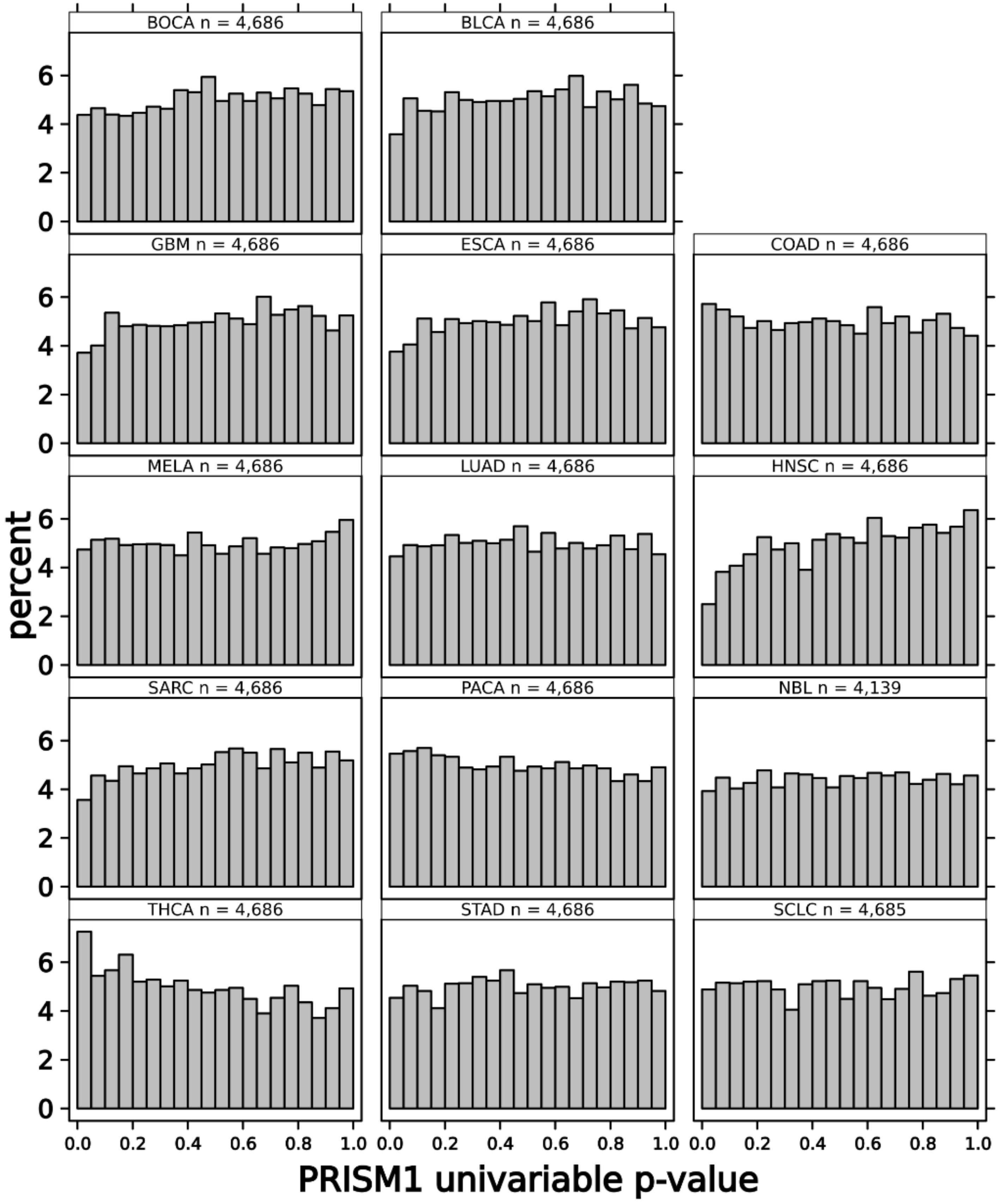

#### Supplementary Figure S9

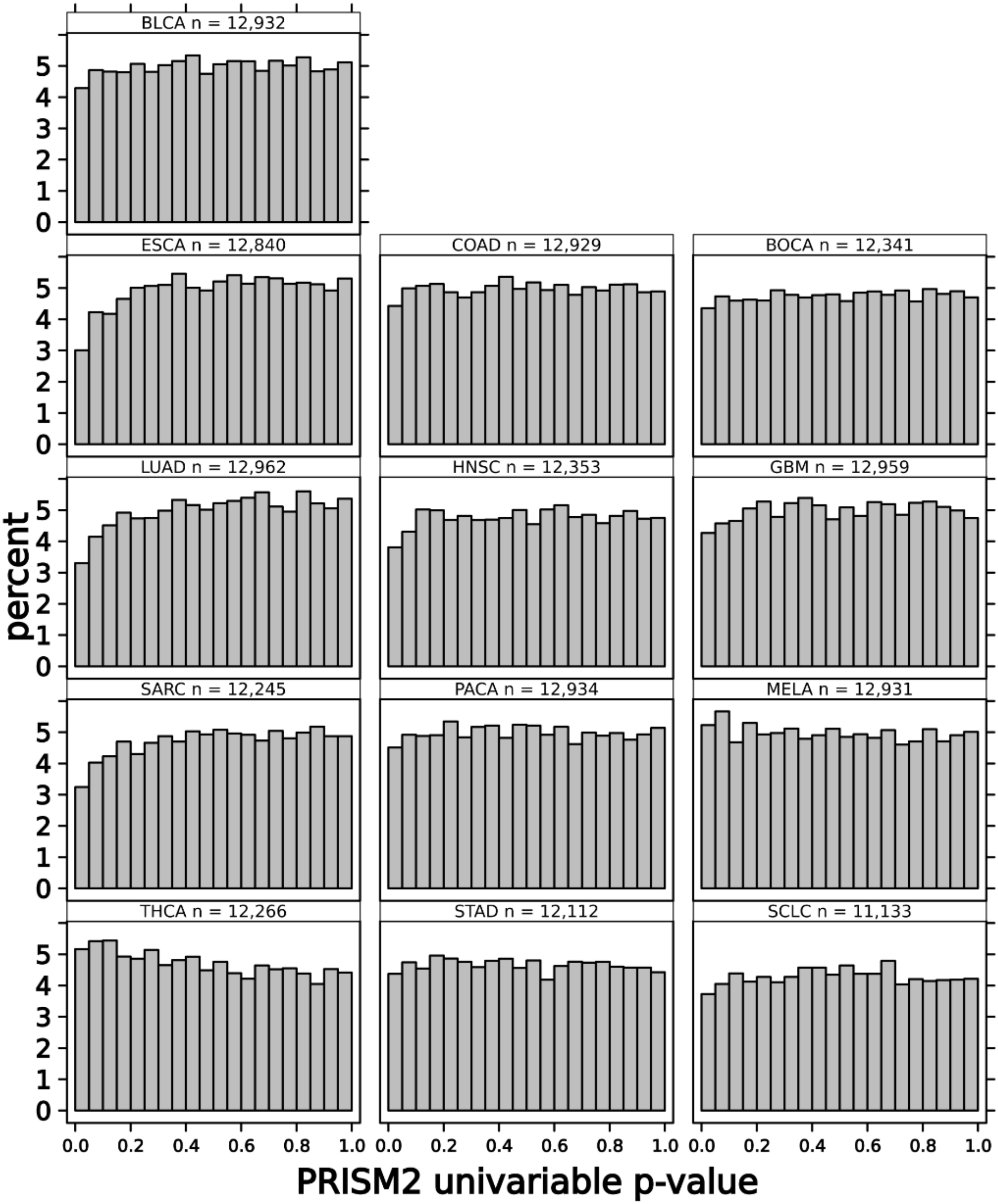
